# *In silico* mouse study identifies tumor growth kinetics as biomarkers for the outcome of anti-angiogenic treatment

**DOI:** 10.1101/300566

**Authors:** Qianhui Wu, Alyssa D. Arnheim, Stacey D. Finley

## Abstract

Angiogenesis is a crucial step in tumor progression, as this process allows tumors to recruit new blood vessels and obtain oxygen and nutrients to sustain growth. Therefore, inhibiting angiogenesis remains a viable strategy for cancer therapy. However, anti-angiogenic therapy has not proved to be effective in reducing tumor growth across a wide range of tumors, and no reliable predictive biomarkers have been found to determine the efficacy of anti-angiogenic treatment. Using our previously established computational model of tumor-bearing mice, we sought to determine whether tumor growth kinetic parameters could be used to predict the outcome of anti-angiogenic treatment. A model trained with datasets from six *in vivo* mice studies was used to generate a randomized *in silico* tumor-bearing mouse population. We analyzed tumor growth in untreated mice (control) and mice treated with an anti-angiogenic agent and determined the Kaplan-Meier survival estimates based on simulated tumor volume data. We found that the ratio between two kinetic parameters, *k*_*0*_ and *k*_*1*_, which characterize the tumor’s exponential and linear growth rates, as well as *k*_*1*_ alone, can be used as prognostic biomarkers of population survival outcome. Our work demonstrates a robust, quantitative approach for identifying tumor growth kinetic parameters as prognostic biomarkers and serves as a template that can be used to identify other biomarkers for anti-angiogenic treatment.

## Background

Tumor angiogenesis results in the vascularization of a tumor. This process facilitates tumor growth by allowing tumor cells to obtain oxygen and nutrients through the newly formed blood vessels. As excessive vascularization is often seen in many types of cancer, inhibiting angiogenesis is thought to decrease tumor growth. Therefore, anti-angiogenic treatment is pursued as an attractive therapeutic strategy in oncology [1, 2].

Bevacizumab is a humanized monoclonal antibody against vascular endothelial growth factor A (VEGF), a key angiogenic promoter in tumors [1]. This drug has been approved as a monotherapy or in combination with chemotherapy for many cancers, including renal cell carcinoma, metastatic colorectal cancer, non-small cell lung cancer, and metastatic cervical cancer [3]. It also gained accelerated approval for treatment of metastatic breast cancer through the US Food and Drug Administration (FDA) in 2008. However, subsequent results showed that bevacizumab failed to improve overall survival and that the drug elicited significant adverse side effects. Consequently, the FDA revoked its approval for use of bevacizumab for first-line metastatic breast cancer in late 2011 [4, 5]. Several Phase II and III clinical stage studies have also revealed contradicting results regarding the benefit of add-on bevacizumab in the neoadjuvant treatment setting for breast cancer patients [6–11]. Altogether, these studies illustrate that angiogenic therapy may not be effective across a wide range of patients. Indeed, breast cancer is a genetically and clinically heterogeneous cancer type, which makes identifying optimal therapies a challenge [12].

More broadly, there is a need for biomarkers to predict the response to treatment and identify the tumors for which anti-angiogenic treatment will be effective. A number of mechanistic biomarkers have been investigated for their ability to predict response to anti-angiogenic treatment and to determine an optimal treatment strategy. Promising biomarker candidates include the concentration ranges of circulating angiogenic molecules (such as plasma levels of VEGF) [13, 14], tissue markers (tumor microvessel density) [15–18], and imaging parameters (MRI-measured *K*^*trans*^) [15, 19, 20]. However, currently no validated and robust biomarkers are available that can guide selection of patients for whom anti-angiogenic therapy is most beneficial [5, 15]].

As an alternative, tumor growth kinetics may be used as biomarkers. There is a body of work that investigates how tumor growth kinetics can serve as prognostic biomarkers of the response to anti-angiogenic treatment [21–25]. Recently, a study showed that volume-based tumor growth kinetics may be a reliable indicator of treatment efficacy, and are in good agreement with standardized approaches for assessing response treatment [21]. Moreover, we developed a computational systems biology model to further investigate the relationship between tumor growth kinetics and the response to anti-angiogenic therapy [26]. The model predicts VEGF distribution and kinetics in tumor-bearing mice, where the dynamic tumor volume is a function of the pro-angiogenic complexes involving VEGF-bound receptors (the “angiogenic signal”). By fitting the model to *in vivo* experimental data, we estimated the kinetic parameters that characterize tumor growth. We then used the trained model to predict the effect of anti-VEGF treatment on tumor volume, using only the estimated parameter values. The model predictions of tumor growth in response to anti-VEGF treatment closely matched experimental data. In this study, we concluded that there is a strong correlation between particular intrinsic kinetic parameters and the response to anti-VEGF treatment in terms of the end relative tumor volume (RTV).

Taking advantage of our established model framework and its strong predictive power, in the present study, we use this model to further investigate the utility of tumor growth kinetics to serve as a biomarker for anti-angiogenic treatment outcome. We performed an *in silico* randomized mouse study and estimated the survival of tumor-bearing mice in response to anti-VEGF treatment. Here, we introduced variability in the mouse population by allowing the tumor growth kinetic parameter values to vary within defined ranges. By generating these large, heterogeneous *in silico* population of tumor-bearing mice, we can eliminate the likely bias caused by animals dropping out in experimental xenograft studies due to limitation of the tumor burden. In general, the average tumor size, particularly in the control group, can be underestimated in an experimental study. This can result in an underestimation of the treatment effect, because large tumors are excluded from the analysis [27]. In contrast, computational modeling avoids these limitations and enables performance metrics (e.g. survival estimates) to be calculated [28]. Furthermore, computational systems biology is a powerful tool for studying how individual components contribute to the function and behavior of a large system, and has been applied to study cancer at multiple scales [29–31]. Such computational models have been used to identify predictive biomarkers and to enhance the efficacy of anti-angiogenic therapies [13, 32, 33].

In our previous work, we focused on the end RTV to evaluate the contribution of parameters to the treatment outcome. In this study, we use more reliable and appropriate readouts. We implemented our tumor growth kinetic model with time-to-event analysis [34]. Specifically, we simulated the Kaplan-Meier survival curves of the *in silico* mice based on the population tumor growth data. We then examined tumor growth kinetic parameters as potential prognostic biomarkers to distinguish the tumor response to anti-angiogenic treatment amongst the stratified groups, by comparing the predicted absolute tumor volume time courses and survival estimates.

## Results

### *In silico* mouse population tumor growth in the whole-body model

We performed an *in silico* randomized mouse study using our whole-body mouse model (**Figure 1**). The model was previously fitted to each of six independent experimental datasets of control tumor volume in mice bearing MDA-MB-231 xenograft tumors and validated with a separate dataset [26]. The values of *k*_*0*_ and *k*_*1*_ (the rates of exponential and linear growth, respectively), *and Ang*_*0*_ (the basal angiogenic signal at time, *t*=0) were estimated. Here, we simulated the tumor growth of the six *in silico* populations of mice (henceforth referred to as “Roland”, “Zibara”, “Tan”, “Volk2008”, “Volk2011a”, and “Volk2011b”), with and without anti-VEGF treatment. For each population, the values of parameters *k*_*0*_ and *k*_*1*_ are randomly varied simultaneously with a uniform distribution within the ranges of their estimated values from our previous model fitting. Previously, a sensitivity analysis showed that the *Ang*_*0*_ parameter was an influential parameter to the model output when the model was fitted; however, further analysis using partial least squares regression (PLSR) indicated that *Ang*_*0*_ was not a strong predictor of response to treatment [26]. Therefore, in each case, *Ang*_*0*_ is set as the median of the range of its estimated values. We generated 400 *in silico* mice for each of the six cases.

**Figure 1.**
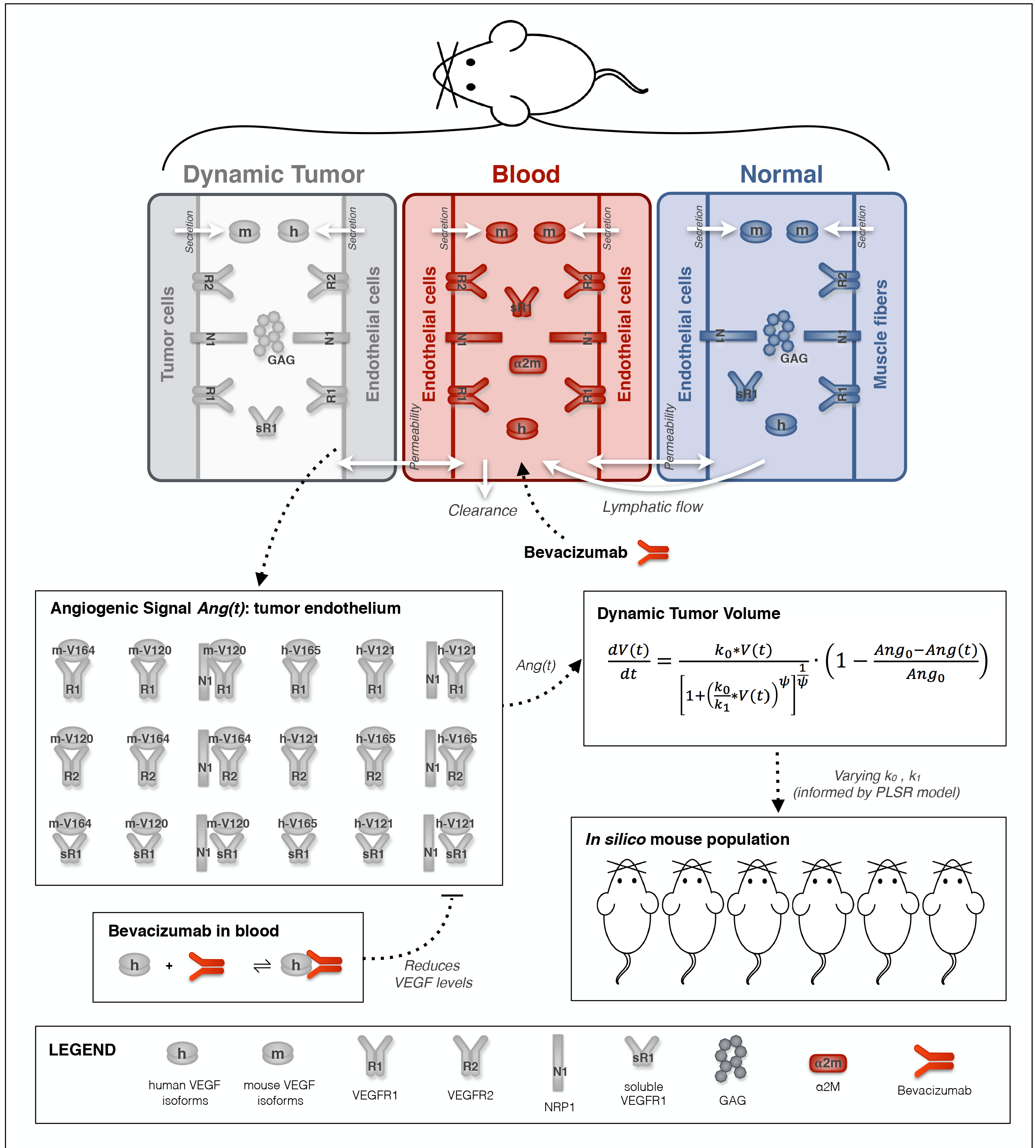
Schematic and overview of computational model of tumor-bearing mice. The three-compartment mouse model was previously trained and validated. The pro-angiogenic signal (*Ang(t)*) is calculated as the summation of the concentrations of VEGF-bound receptor complexes in the tumor endothelium. The dynamic tumor volume is a function of the angiogenic signal. In this study, we randomly varied tumor growth parameters within specified ranges to simulated tumor growth of several heterogeneous mouse populations.

Our simulations show that among the six cases, the anti-VEGF treatment has differential effects in reducing the tumor growth, as compared to the control group (**Figure 2**). For all cases, we used a single treatment protocol different from protocols used in each of the six experimental studies, in order to compare the predicted results without bias (termed “protocol A”). For Roland, Tan, Volk2008, and Volk2011b (**Figure 2A,C,D,F**), the treated tumor volumes are less than the untreated tumors. Meanwhile, for Zibara and Volk2011a (**Figure 2B,E**), there is no apparent difference in the tumor volumes for the treated and control groups. Thus, the model simulations reveal distinct differences in the effect of anti-VEGF treatment.

**Figure 2.**
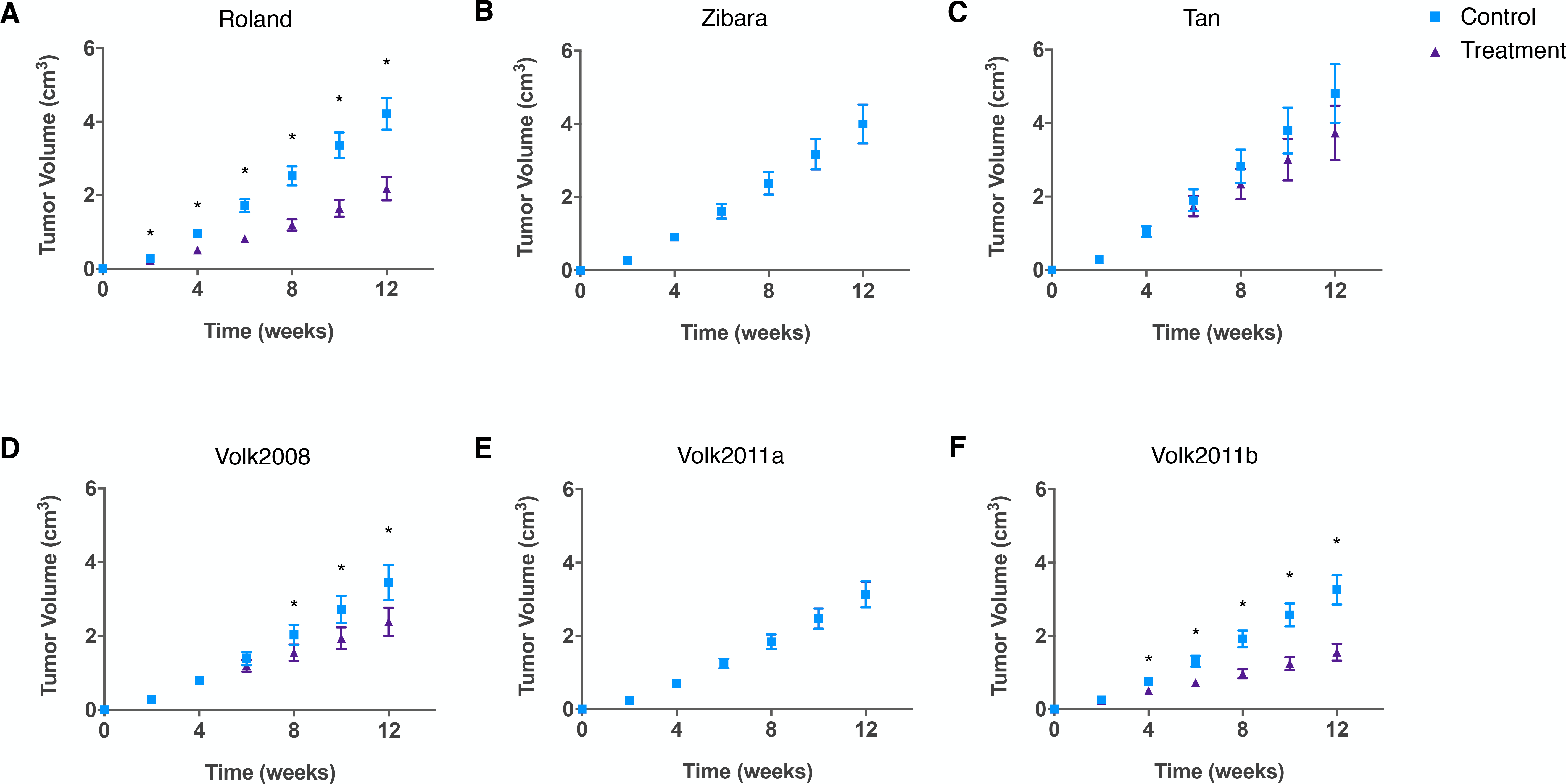
Model-simulated tumor growth data of *in silico* mouse populations. The whole-body mouse model previously fit to each of the six datasets individually was used to simulate tumor volume over time. To generate the simulated tumors, the tumor growth kinetic parameters *k*_0_ and *k_1_* were randomly varied within the range of the estimated values. A total of 400 simulations were run for each case. The mean and 95% confidence interval at each time point are shown. **A**, Roland. **B**, Zibara. **C**, Tan. **D**, Volk2008. **E**, Volk2011a. **F**, Volk2011b. Asterisks indicate that the difference between the control and treatment group tumor volumes is statistically significant (*p*<0.05).

We further studied the effect of anti-VEGF treatment on tumor growth using RTV, the ratio between the mean tumor volumes of the treated and control groups. We calculated the RTV at each time point for all simulated tumors (**Figure S1**). We also determined the RTV at the end of treatment (**Figure S2**). The RTV values in all cases are smaller than one, indicating that the anti-VEGF treatment limits tumor growth, similar to what has been observed experimentally [35–39]. For Zibara and Volk2011a, the endpoint RTV values are just slightly less than one (**Figure S2B,E**), which is an expected result based on the similar tumor growth curves between the control and treated groups (**Figure 2B,E**). Comparing the endpoint RTV among all six cases, the effect of anti-VEGF treatment in limiting tumor growth is the strongest for Volk2011b (RTV = 0.459 ± 0.054), followed by Roland (0.454 ± 0.096), Volk2008 (0.615 ± 0.066), and Tan (0.638 ± 0.049). This treatment effect is the least significant in Zibara (0.979 ± 0.009) and Volk2011a (0.987 ± 0.013).

### Kinetic parameters as potential predictor for stratified population response

We investigated the relationship between the parameters that characterize tumor growth kinetics and the effect of the anti-VEGF treatment. Previously, our PLSR analysis indicated that for nearly all pairwise comparisons, if the RTV values for two datasets were significantly different, their *k*_*0*_/*k*_*1*_ ratios were also significantly different. This implies that the *k*_*0*_/*k*_*1*_ is a large contributor in predicting the endpoint RTV [26]. Additionally, plotting the RTV versus *k*_*0*_, *k*_*1*_, and *k*_*0*_/*k*_*1*_ shows some relationship between the endpoint RTV and the tumor growth parameters (**Figure S2**). Therefore, we investigated whether these tumor growth parameters could stratify the simulated mouse populations, and distinguish their tumor growth and survival estimates. To address this question, we used our simulated tumor growth data for each case, noting the number of *in silico* mice at each time point. We record the time at which a mouse is “sacrificed”, which happens when the tumor volume reaches 2 cm^3^, as typically done in experimental studies [40]. This approach for modeling population survival allows us to closely mimic the practice in preclinical animal studies, and provides easily interpretable insights for researchers and clinicians.

We used the simulated population survival data to determine if *k*_*0*_, *k*_*1*_, or *k*_*0*_/*k*_*1*_ can be used to discriminate between tumors for which anti-VEGF treatment is effective or not. We found that in each case, a range of *k*_*0*_/*k*_*1*_ ratios, as well as *k*_*1*_, can be used to distinguish the population response to the anti-VEGF treatment (**Figure 3B,C**). We term these “*ratio_thresh_”* and “*k*_*1,thresh*_”, the values of the growth kinetic parameters that separate the simulated mouse population into groups with significantly different survival estimates. In contrast, we did not find any values of *k*_*0*_ alone that could be used to separate the simulated mouse population into groups whose survival estimates are statistically different for Roland, Zibara, and Volk2011b cases. For Tan and Volk2008, we only found one such *k*_*0*_ value in each case (**Figure 3A**).

**Figure 3.**
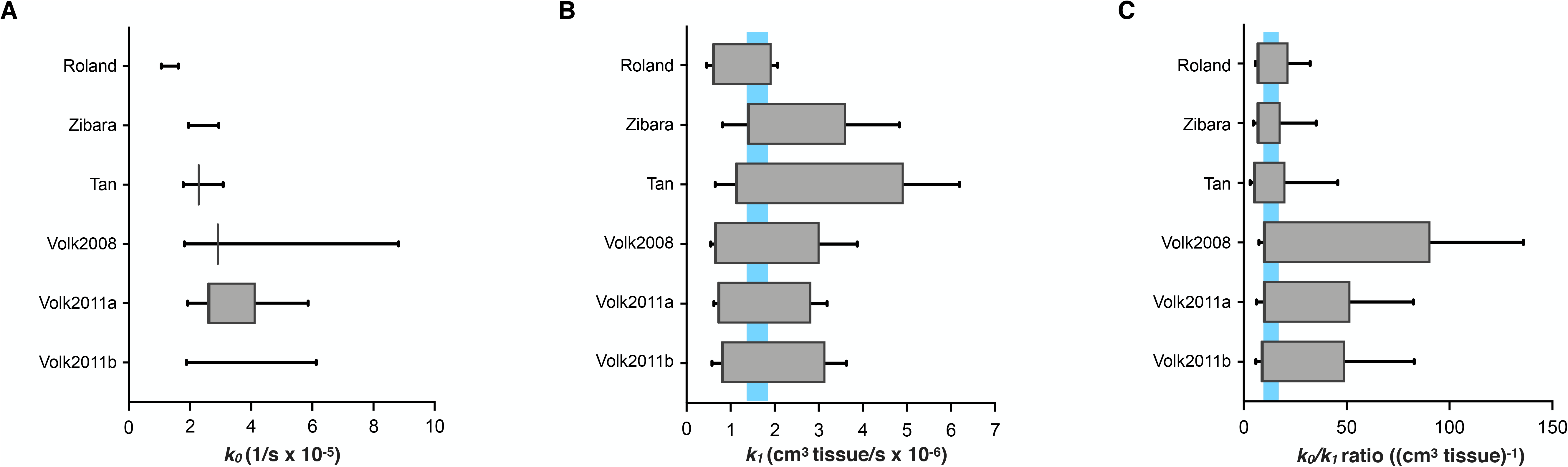
Range of parameter and threshold values. In each of the six cases, values of *k*_*1,thresh*_ and *ratio*_*thresh*_ were found among all of the randomly generated values of *k*_*1*_ or *k*_*0*_/*k*_*1*_ ratio used in the simulations. **A**, *k*_*0*_. **B**, *k_1_.* **C**, *k*_*0*_/*k*_*1*_ ratio. *Bars:* the ranges of all generated parameter values in each case. *Boxes:* the ranges of possible threshold values in each case. *Shading:* the common range of threshold values among the six cases.

Interestingly, although the ranges of generated *k*_*0*_/*k*_*1*_ ratios and *k*_*1*_ were different for each of the six sets of tumor growth data, we found that there is an overlap among the potential *ratio_thresh_* or *k*_1,thresh_ values found in each of the six cases. The common range of *ratio_thresh_* is 9.757 to 17.982, and that of *k*_*1,thresh*_ is 1.391 ×10^−6^ to 1.931 ×10^−6^. This means that separating the treatment group by any *k*_*1,thresh*_ or *ratio*_*thresh*_ value within its respective range will produce two groups of treated mice that have statistically different survival estimates. Specifically, the treated group with *k*_*0*_/*k*_*1*_ ratios larger than the *ratio*_*thresh*_ value has a better survival estimate than the treated group with smaller ratios. The treated group with *k*_*1*_ smaller than the *k*_*1,thresh*_ value has a better survival estimate than the treated group with larger *k*_*1*_.

We used the median *ratio*_*thresh*_ value to illustrate this distinction. We compare the survival estimates for a total of six groups: 1) all mice in the control group; 2) all mice in the treatment group; 3) control group with *k*_*0*_/*k*_*1*_ < *ratio*_*thresh*_; 4) control group with *k*_*0*_/*k*_*1*_ > *ratio*_*thresh*_; 5) treatment group with *k*_*0*_/*k*_*1*_ < *ratio*_*thresh*_; and 6) treatment group with *k*_*0*_/*k*_*1*_ > *ratio*_*thresh*_. We generated the Kaplan-Meier survival curves for these groups for each of the six cases investigated (**Figure 4**). We also estimated the median survival of the six groups in each case (**Table 1**), the Mantel-Haenszel hazard ratio (HR), with 95% confidence interval (CI), and the *p*-values from the Mantel-Cox log rank test for the survival curve comparison (**Table 2**). When comparing two groups, if the HR is less than one, the first group has a lower death rate (see Methods). Together these analyses emphasize that mice with larger *k*_*0*_/*k*_*1*_ ratios survive for longer, with *p*-value < 0.05. Interestingly, for Zibara and Volk2011a, although the anti-VEGF treatment does not significantly reduce tumor growth and therefore does not yield a better survival estimate for the treated groups compared to their control groups (**Figures 2B,E** **and 4B,E**), the stratified groups yield significantly different survival estimates. That is, the control and treated groups with *k*_*0*_/*k*_*1*_ ratios larger than *ratio*_*thresh*_ have better survival estimates than those with smaller *k*_*0*_/*k*_*1*_ ratios.

**Figure 4.**
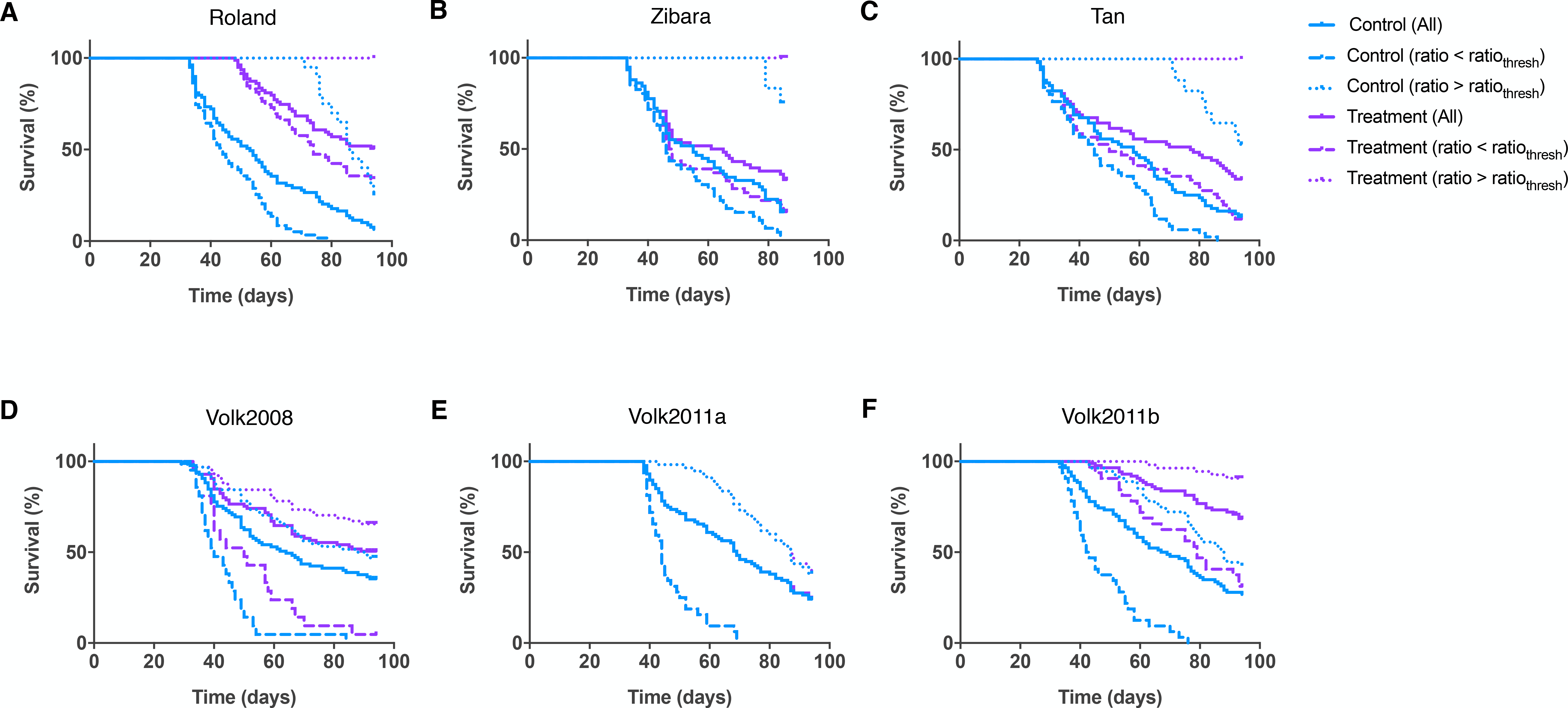
Kaplan-Meier curves for the six simulated groups of tumor-bearing mice. Here, the *ratio*_*thresh*_ value is taken as the median from the common range found among the six cases. **A**, Roland. **B**, Zibara. **C**, Tan. **D**, Volk2008. **E**, Volk2011a. **F**, Volk2011b. Theestimated survival curves of *in silico* mice subgroups within each group are shown in each plot: all mice, mice with ratio above or below the median *ratio_thresh_* in the control setting or with treatment.

**Table 1.** Summary of median survival of population separated by median *ratio*_*thresh*_

| Median survival (days) | Roland<sup>†</sup> | Zibara<sup>†</sup> | Tan<sup>†</sup> | Volk 2008<sup>†</sup> | Volk 2011a<sup>†</sup> | Volk 2011b<sup>†</sup> | Zibara<sup>§</sup> | Volk 2011a<sup>¶</sup> | Volk 2011a<sup>¶¶</sup> | Mollard<sup>†</sup> |
| --- | --- | --- | --- | --- | --- | --- | --- | --- | --- | --- |
| Control (All) | 53 | 55 | 58 | 64 | 69 | 68 | 55 | 69 | 69 | 85 |
| Control ($k_0/k_1 < ratio_{thresh}$) | 44 | 44.5 | 45 | 40 | 44 | 42.5 | 44.5 | 42.5 | 44 | 77 |
| Control ($k_0/k_1 > ratio_{thresh}$) | 86.5 | 84 | Un* | 89.5 | 87 | 88 | 84 | 80 | 87 | 96 |
| Treatment (All) | Un* | 63 | 77.5 | Un* | 69 | Un* | 63 | 71 | Un* | Un* |
| Treatment ($k_0/k_1 < ratio_{thresh}$) | 74 | 46 | 50 | 50 | 44 | 79 | 46 | 42.5 | 78.5 | Un* |
| Treatment ($k_0/k_1 > ratio_{thresh}$) | Un* | Un* | Un* | Un* | 87 | Un* | Un* | 87 | Un* | Un* |

\*Un: Undefined. The median survival cannot be estimated if the survival estimation does not reach below 50%;
<sup>†</sup>protocol A: biweekly treatment at a dosage of 10 mg/kg, starting when tumor volume reaches 0.1 cm<sup>3</sup>;
<sup>§</sup>protocol Z: biweekly treatment at a dosage of 10 mg/kg, starting when tumor volume is 0.004 cm<sup>3</sup> (upon engraftment of tumor);
<sup>¶</sup>protocol V11a: biweekly treatment at a dosage of 10 mg/kg, starting when tumor volume reaches 0.5 cm<sup>3</sup>;
<sup>¶¶</sup>protocol V11a-D: biweekly treatment at a dosage of 20 mg/kg, starting when tumor volume reaches 0.5 cm<sup>3</sup>.

**Table 2.** Statistics comparing the Kaplan-Meier survival 1 curves of population separated by median *ratio*_*thresh*_: hazard ratio (95%CI) and log rank test *p*-values

| HR<br>(95% CI)<br>p-value | Roland<sup>†</sup> | Zibara<sup>‡</sup> | Tan<sup>†</sup> | Volk2008<sup>†</sup> | Volk2011a<sup>†</sup> |
| --- | --- | --- | --- | --- | --- |
| Treatment<br>($k_0/k_1 > ratio_{thresh}$) vs.<br>Treatment<br>($k_0/k_1 < ratio_{thresh}$) | 0.2073<br>(0.1059-0.4057)<br>p<0.0001 | 0.2005<br>(0.0991-0.4055)<br>p<0.0001 | 0.1623<br>(0.0872-0.3021)<br>p<0.0001 | 0.0576<br>(0.0237-0.1399)<br>p<0.0001 | 0.0216<br>(0.0098-0.0476)<br>p<0.0001 |
| Control<br>($k_0/k_1 > ratio_{thresh}$) vs.<br>Control<br>($k_0/k_1 < ratio_{thresh}$) | 0.1214<br>(0.06974-0.2144)<br>p<0.0001 | 0.1627<br>(0.0866-0.3056)<br>p<0.0001 | 0.1445<br>(0.0808-0.2582)<br>p<0.0001 | 0.0422<br>(0.0173-0.1035)<br>p<0.0001 | 0.0296<br>(0.0098-0.0476)<br>p<0.0001 |
| Treatment<br>($k_0/k_1 > ratio_{thresh}$) vs.<br>Control<br>($k_0/k_1 > ratio_{thresh}$) | 0.0675<br>(0.0234-0.1949)<br>p<0.0001 | 0.1191<br>(0.01194-1.188)<br>p=0.0697 | 0.101<br>(0.0249-0.4103)<br>p=0.0013 | 0.5683<br>(0.33490-9643)<br>p=0.0362 | 0.9921<br>(0.6117-1.609)<br>p=0.9742 |
| Treatment<br>($k_0/k_1 > ratio_{thresh}$) vs.<br>Treatment (All) | 0.2562<br>(0.1239-0.5299)<br>p=0.0002 | 0.2538<br>(0.1191-0.5408)<br>p=0.0004 | 0.239<br>(0.1236-0.4621)<br>p<0.0001 | 0.6138<br>(0.3732-1.01)<br>p=0.0546 | 0.576<br>(0.3863-0.8588)<br>p=0.0068 |
| Treatment (All) vs.<br>Control (All) | 0.2307<br>(0.1548-0.3438)<br>p<0.0001 | 0.6742<br>(0.4384-1.037)<br>p=0.0726 | 0.5845<br>(0.3934-0.8684)<br>p=0.0079 | 0.6481<br>(0.4331-0.9699)<br>p=0.0350 | 0.9959<br>(0.7038-1.409)<br>p=0.9815 |
| Treatment<br>($k_0/k_1 < ratio_{thresh}$) vs.<br>Treatment (All) | 1.569<br>(0.9721-2.41)<br>p=0.0549 | 1.558<br>(0.9714-2.5)<br>p=0.0658 | 1.794<br>(1.158-2.778)<br>p=0.0089 | 6.405<br>(2.971-13.81)<br>p<0.0001 | 7.657<br>(4.065-14.42)<br>p<0.0001 |
| Treatment<br>($k_0/k_1 > ratio_{thresh}$) vs.<br>Treatment<br>($k_0/k_1 < ratio_{thresh}$) | 0.0673<br>(0.0288-0.157)<br>p<0.0001 | 0.2332<br>(0.1191-0.4566)<br>p<0.0001 | 0.01104<br>(0.0039-0.0315)<br>p<0.0001 | 0.358<br>(0.0157-0.0815)<br>p<0.0001 | 0.1862<br>(0.1043-0.3327)<br>p<0.0001 |
| Control<br>($k_0/k_1 > ratio_{thresh}$) vs.<br>Control<br>($k_0/k_1 < ratio_{thresh}$) | 0.0368<br>(0.01750-0.0778)<br>p<0.0001 | 0.1655<br>(0.0925-0.2962)<br>p<0.0001 | 0.0110<br>(0.0039-0.0315)<br>p<0.0001 | 0.0418<br>(0.0204-0.0857)<br>p<0.0001 | 0.6573<br>(0.5219-0.8029)<br>p<0.0001 |
| Treatment<br>($k_0/k_1 > ratio_{thresh}$) vs.<br>Control<br>($k_0/k_1 > ratio_{thresh}$) | 0.1592<br>(0.0817-0.3102)<br>p<0.0001 | 0.1707<br>(0.0557-0.5226)<br>p=0.002 | 0.7027<br>(0.4718-1.047)<br>p=0.0825 | 0.0842<br>(0.0452-0.1568)<br>p<0.0001 | 0.0782<br>(0.0561-0.1089)<br>p<0.0001 |
| Treatment<br>($k_0/k_1 > ratio_{thresh}$) vs.<br>Treatment (All) | 0.3364<br>(0.1662-0.6806)<br>p=0.0024 | 0.2513<br>(0.1294-0.4881)<br>p<0.0001 | 0.7306<br>(0.5008-1.066)<br>p=0.1033 | 0.2524<br>(0.126-0.5058)<br>p=0.0001 | 0.317<br>(0.176-0.571)<br>p=0.0001 |
| Treatment (All) vs.<br>Control (All) | 0.266<br>(0.1733-0.4083)<br>p<0.0001 | 0.6742<br>(0.4384-1.037)<br>p=0.0726 | 0.7766<br>(0.5543-1.088)<br>p=0.1416 | 0.2016<br>(0.1343-0.3027)<br>p<0.0001 | 0.0940<br>(0.0762-0.1159)<br>p<0.0001 |
| Treatment<br>($k_0/k_1 < ratio_{thresh}$) vs.<br>Treatment (All) | 3.938<br>(1.985-7.811)<br>p<0.0001 | 2.119<br>(1.294-3.47)<br>p=0.0005 | 15.64<br>(6.536-37.41)<br>p<0.0001 | 5.074<br>(2.653-9.705)<br>p<0.0001 | 0.5152<br>(0.3481-0.8458)<br>p=0.0035 |

<sup>†</sup>protocol A, <sup>§</sup>protocol Z, <sup>¶</sup>protocol V11a, <sup>¶¶</sup>protocol V11a-D

We performed a similar analysis using the median *k*_*1,thresh*_ value to show the distinction between the survival estimates (**Figure S3**). The control and treated groups with *k*_*1*_ smaller than *k*_*1, thresh*_ have better survival estimates than those with larger *k*_*1*_ values. We also estimated the median survival of the six groups separated using the median *k*_*1,thresh*_ (**Table 3**), the Mantel-Haenszel HR, and the *p*-values from the Mantel-Cox log rank test for the survival curve comparison (**Table 4**). From these analyses, mice with smaller *k*_*1*_ survive longer than those with larger *k*_*1*_, and the HR is smaller than one (*p*<0.05).

**Table 3.** Summary of median survival of population separated by median *k*_*1,thresh*_

| Median survival<br>(days) | Roland<sup>†</sup> | Zibara<sup>†</sup> | Tan<sup>†</sup> | Volk<br>2008<sup>†</sup> | Volk<br>2011a<sup>†</sup> | Volk<br>2011b<sup>†</sup> | Zibara<sup>§</sup> | Volk<br>2011a<sup>¶</sup> | Volk<br>2011a<sup>¶¶</sup> | Mollard<sup>†</sup> |
| --- | --- | --- | --- | --- | --- | --- | --- | --- | --- | --- |
| Control (All) | 53 | 55 | 58 | 64 | 69 | 68 | 55 | 69 | 69 | 87 |
| Control<br>($k_1 < k_{1,thresh}$) | 59.5 | 84 | 86 | Un* | 87 | Un* | 84 | 87 | 87 | 92 |
| Control<br>($k_1 > k_{1,thresh}$) | 35 | 45 | 43 | 41 | 44 | 45 | 45 | 44 | 44 | 72 |
| Treatment<br>(All) | Un* | 63 | 77.5 | Un* | 69 | Un* | 63 | 71 | Un* | Un* |
| Treatment<br>($k_1 < k_{1,thresh}$) | Un* | Un* | Un* | Un* | 88 | Un* | 108 | 108 | Un* | Un* |
| Treatment<br>($k_1 > k_{1,thresh}$) | 55 | 46 | 46 | 47.5 | 44 | 82 | 44 | 44 | 78.5 | Un* |

<sup>†</sup>protocol A, <sup>§</sup>protocol Z, <sup>¶</sup>protocol V11a, <sup>¶¶</sup>protocol V11a-D

**Table 4.**
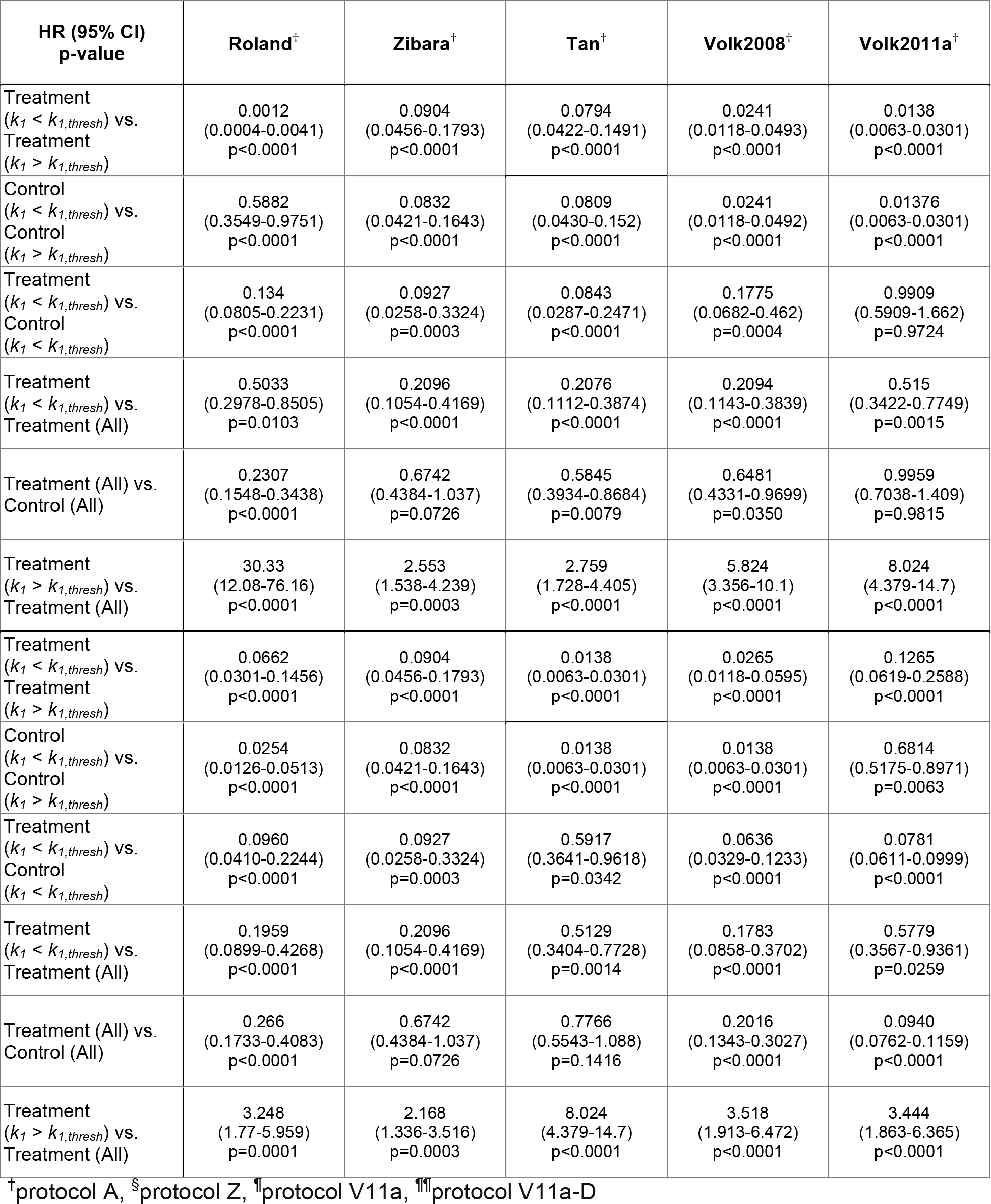
Statistics comparing the Kaplan-Meier survival curves of population 9 separated by median *k*_*1,thresh*_: hazard ratio (95%CI) and log rank test *p*-values

| HR (95% CI)<br>p-value | Roland<sup>†</sup> | Zibara<sup>‡</sup> | Tan<sup>§</sup> | Volk2008<sup>†</sup> | Volk2011a<sup>†</sup> |
| --- | --- | --- | --- | --- | --- |
| Treatment<br>($k_1 < k_{1,thresh}$) vs.<br>Treatment<br>($k_1 > k_{1,thresh}$) | 0.0012<br>(0.0004-0.0041)<br>p<0.0001 | 0.0904<br>(0.0456-0.1793)<br>p<0.0001 | 0.0794<br>(0.0422-0.1491)<br>p<0.0001 | 0.0241<br>(0.0118-0.0493)<br>p<0.0001 | 0.0138<br>(0.0063-0.0301)<br>p<0.0001 |
| Control<br>($k_1 < k_{1,thresh}$) vs.<br>Control<br>($k_1 > k_{1,thresh}$) | 0.5882<br>(0.3549-0.9751)<br>p<0.0001 | 0.0832<br>(0.0421-0.1643)<br>p<0.0001 | 0.0809<br>(0.0430-0.152)<br>p<0.0001 | 0.0241<br>(0.0118-0.0492)<br>p<0.0001 | 0.01376<br>(0.0063-0.0301)<br>p<0.0001 |
| Treatment<br>($k_1 < k_{1,thresh}$) vs.<br>Control<br>($k_1 < k_{1,thresh}$) | 0.134<br>(0.0805-0.2231)<br>p<0.0001 | 0.0927<br>(0.0258-0.3324)<br>p=0.0003 | 0.0843<br>(0.0287-0.2471)<br>p<0.0001 | 0.1775<br>(0.0682-0.462)<br>p=0.0004 | 0.9909<br>(0.5909-1.662)<br>p=0.9724 |
| Treatment<br>($k_1 < k_{1,thresh}$) vs.<br>Treatment (All) | 0.5033<br>(0.2978-0.8505)<br>p=0.0103 | 0.2096<br>(0.1054-0.4169)<br>p<0.0001 | 0.2076<br>(0.1112-0.3874)<br>p<0.0001 | 0.2094<br>(0.1143-0.3839)<br>p<0.0001 | 0.515<br>(0.3422-0.7749)<br>p=0.0015 |
| Treatment (All) vs.<br>Control (All) | 0.2307<br>(0.1548-0.3438)<br>p<0.0001 | 0.6742<br>(0.4384-1.037)<br>p=0.0726 | 0.5845<br>(0.3934-0.8684)<br>p=0.0079 | 0.6481<br>(0.4331-0.9699)<br>p=0.0350 | 0.9959<br>(0.7038-1.409)<br>p=0.9815 |
| Treatment<br>($k_1 > k_{1,thresh}$) vs.<br>Treatment (All) | 30.33<br>(12.08-76.16)<br>p<0.0001 | 2.553<br>(1.538-4.239)<br>p=0.0003 | 2.759<br>(1.728-4.405)<br>p<0.0001 | 5.824<br>(3.356-10.1)<br>p<0.0001 | 8.024<br>(4.379-14.7)<br>p<0.0001 |
| Treatment<br>($k_1 < k_{1,thresh}$) vs.<br>Treatment<br>($k_1 > k_{1,thresh}$) | 0.0662<br>(0.0301-0.1456)<br>p<0.0001 | 0.0904<br>(0.0456-0.1793)<br>p<0.0001 | 0.0138<br>(0.0063-0.0301)<br>p<0.0001 | 0.0265<br>(0.0118-0.0595)<br>p<0.0001 | 0.1265<br>(0.0619-0.2588)<br>p<0.0001 |
| Control<br>($k_1 < k_{1,thresh}$) vs.<br>Control<br>($k_1 > k_{1,thresh}$) | 0.0254<br>(0.0126-0.0513)<br>p<0.0001 | 0.0832<br>(0.0421-0.1643)<br>p<0.0001 | 0.0138<br>(0.0063-0.0301)<br>p<0.0001 | 0.0138<br>(0.0063-0.0301)<br>p<0.0001 | 0.6814<br>(0.5175-0.8971)<br>p=0.0063 |
| Treatment<br>($k_1 < k_{1,thresh}$) vs.<br>Control<br>($k_1 < k_{1,thresh}$) | 0.0960<br>(0.0410-0.2244)<br>p<0.0001 | 0.0927<br>(0.0258-0.3324)<br>p=0.0003 | 0.5917<br>(0.3641-0.9618)<br>p=0.0342 | 0.0636<br>(0.0329-0.1233)<br>p<0.0001 | 0.0781<br>(0.0611-0.0999)<br>p<0.0001 |
| Treatment<br>($k_1 < k_{1,thresh}$) vs.<br>Treatment (All) | 0.1959<br>(0.0899-0.4268)<br>p<0.0001 | 0.2096<br>(0.1054-0.4169)<br>p<0.0001 | 0.5129<br>(0.3404-0.7728)<br>p=0.0014 | 0.1783<br>(0.0858-0.3702)<br>p<0.0001 | 0.5779<br>(0.3567-0.9361)<br>p=0.0259 |
| Treatment (All) vs.<br>Control (All) | 0.266<br>(0.1733-0.4083)<br>p<0.0001 | 0.6742<br>(0.4384-1.037)<br>p=0.0726 | 0.7766<br>(0.5543-1.088)<br>p=0.1416 | 0.2016<br>(0.1343-0.3027)<br>p<0.0001 | 0.0940<br>(0.0762-0.1159)<br>p<0.0001 |
| Treatment<br>($k_1 > k_{1,thresh}$) vs.<br>Treatment (All) | 3.248<br>(1.77-5.959)<br>p=0.0001 | 2.168<br>(1.336-3.516)<br>p=0.0003 | 8.024<br>(4.379-14.7)<br>p<0.0001 | 3.518<br>(1.913-6.472)<br>p<0.0001 | 3.444<br>(1.863-6.365)<br>p<0.0001 |

<sup>†</sup>protocol A, <sup>§</sup>protocol Z, <sup>‡</sup>protocol V11a, <sup>¶</sup>protocol V11a-D

### Alternative treatment strategies to improve survival estimates

We next sought to understand whether alternative treatment protocols can effectively reduce tumor volume for the Zibara and Volk2011a cases, since the baseline protocol we used did not significantly affect tumor volume. For the Zibara case, we simulated the original treatment protocol used in the experimental study (termed “protocol Z”). This protocol starts the 10 mg/kg biweekly treatment upon tumor engraftment (assuming the initial tumor volume to be 0.004 cm^3^) [36]. The predicted tumor volumes are smaller in the treated group (**Figure S4A**), recapitulating the findings from the published experimental study. The predictions may suggest that in this case, starting the treatment earlier is more effective in limiting the tumor growth. For mice with *k*_*0*_/*k*_*1*_ ratios larger than the median *ratio*_*thresh*_ value (13.689), or with *k*_*1*_ smaller than the median *k*_*1,thresh*_ value (1.661 × 10^−6^), the HR between the treated and control groups is smaller than one, and the survival curves are significantly different (*p*<0.0001) (**Tables 2** **and 4**).

For Volk2011a, we simulated treatment termed “protocol V11a”, which starts the 10 mg/kg biweekly treatment when the tumor volume reaches 0.5 cm^3^, a start time extracted from the published preclinical study [39]. After 12 weeks, the simulated mean tumor volumes in the treated group are significantly smaller than the control tumors (**Figure S4B**). However, the survival estimates were not significantly different (*p*>0.05). Again, the treated group with *k*_*0*_/*k*_*1*_ ratios larger than the median *ratio*_*thresh*_, or with *k*_*1*_ smaller than the median *k*_*1,thresh*_, has a significantly better survival estimate than the opposite group (*p*<0.0001) (**Tables 2** **and 4**). This phenomenon is similar to that observed in the Volk2011a case using protocol A, where the two groups separated according to the *k*_*0*_/*k*_*1*_ ratio or *k*_*1*_ have distinct survival estimates, but there is no significant difference between the treated and control groups.

Finally, we explored whether another treatment protocol could significantly improve the survival estimates for the treated group compared to the control. We simulated protocol V11a-D, where biweekly treatment starts when the tumor volume reaches 0.5 cm^3^, and the drug dosage is doubled to 20 mg/kg. This treatment protocol significantly limits the tumor growth (**Figure S4C**), and the survival curves are significantly better for the treated group compared to the control (*p*<0.0001). Overall, the treated and control groups have an HR of 0.2016 (95% CI: 0.1343-0.3027) (**Table 2**).

### Validation of thresholds using an independent dataset

To validate the use of the range of *ratio*_*thresh*_ and *k_1,thresh_* values that we found, we used a recently published independent set of data that measures tumor growth in mice with MDA-MB-231 xenografts, with or without bevacizumab treatment [41]. First, we fit the model to the measured tumor volumes without treatment. We obtained 12 sets of estimated parameter values for *k*_*0*_, *k*_*1*_, and *Ang*_*0*_ that allow the model to best fit to the control data. We then validated the fitted model by simulating anti-VEGF treatment and comparing to the experimental measurements. The predicted tumor growth with treatment matches closely to the experimental data (**Figure 5A**).

**Figure 5.**
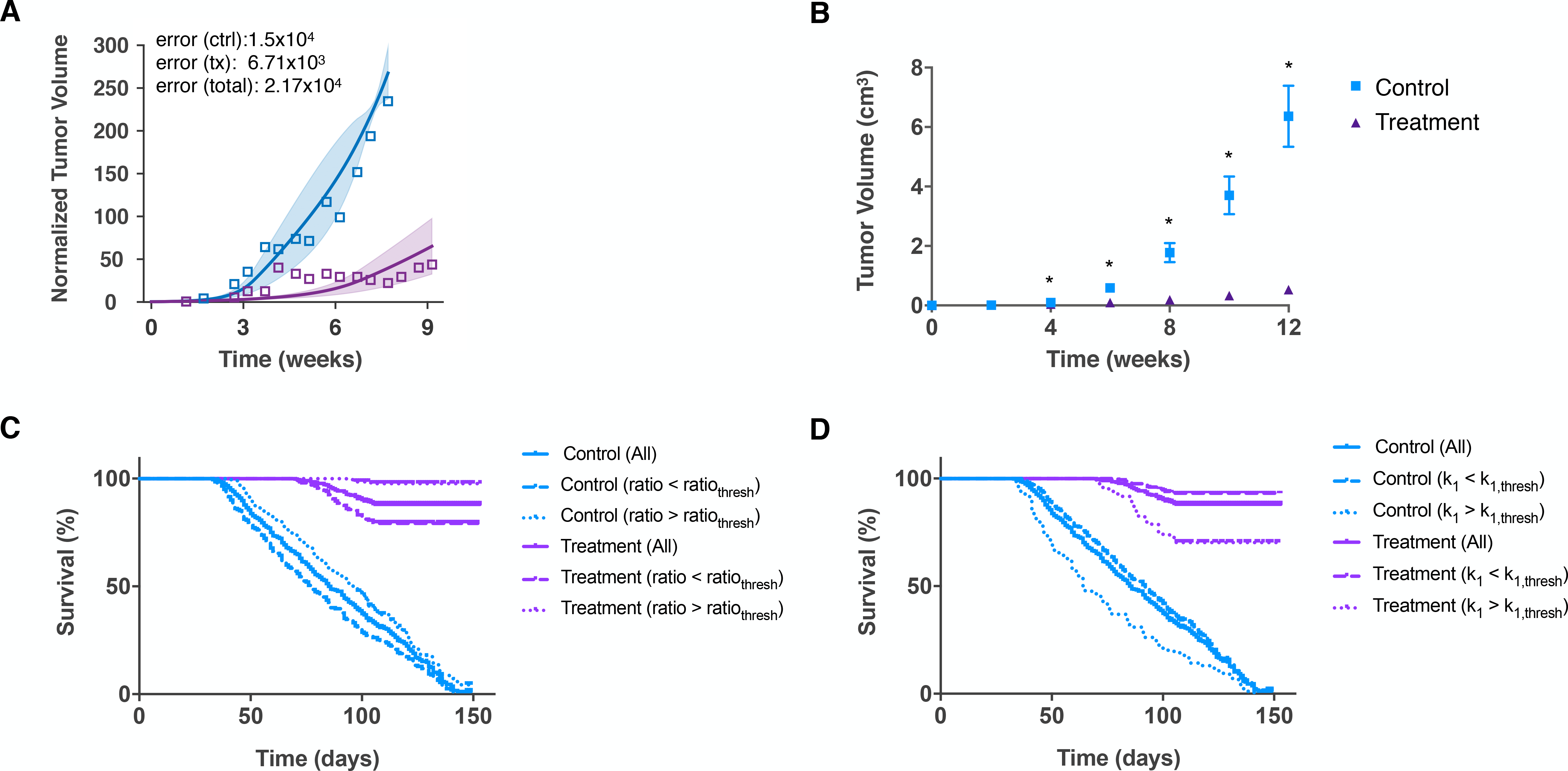
Validation of *ratio*_*thresh*_ and *k*_*1,thresh*_ values with an independent set of data from [41]. **A**, Model fit to control data and validation with treatment data from [41]. The model was fit to normalized tumor volume, and the tumor growth kinetic parameters were estimated. The model is able to reproduce experimental data in the control group and predict the treatment data. *Line:* mean of best fits. *Shading:* range of standard deviation. *Squares:* experimental data. *Error values:* SSR for mean of the best fits. **B**, Model-simulated tumor growth of an *in silico* mouse population, with tumor growth kinetic parameters *k*_*0*_ and *k*_*1*_ for each simulation randomly varied within the range of their estimated values. The mean and 95% confidence interval at each time point are shown. Asterisks indicate that the difference between the control and treatment group tumor volumes is statistically significant (*p*<0.05). **C and D**, Estimated Kaplan-Meier survival curves of the simulated mouse population obtained using the model that was fitted to Mollard data. The population is separated using the median of the range of **C**, *ratio*_*thresh*_ values (13.8693), or **D**, *k*_*1,thresh*_ values (1.661 × 10^−6^).

Using the same approach as described above, we generated 400 sets of tumor volumes for an *in silico* mouse population with and without treatment (referred to as “Mollard”). To do so, we randomly varied *k*_*0*_ and *k*_*1*_ from the ranges of the 12 sets of estimated parameter values from model fitting to the Mollard dataset, with *Ang*_*0*_ held constant at the median of its estimated values. The simulated tumor volumes for the control and treated groups are shown in **Figure 5B**.

We generated the population survival data based on the simulated tumor growth profiles. We tested whether the common range of *ratio*_*thresh*_ and *k*_*1,thresh*_ values identified using the six datasets described above are able to separate the population survival data for this validation case (Mollard). For all *ratio*_*thresh*_ values within the range, the survival estimate of the treated mice with *k*_*0*_/*k*_*1*_ ratios larger than the threshold is better than those with smaller *k*_*0*_/*k*_*1*_ ratios. Examples using the median *ratio*_*thresh*_ and the median *k*_*1,thresh*_ are shown in **Figure 5C-D**. We calculated the HR values, as well as the *p*-value from the Mantel-Cox log rank test among the treated and control groups, separated using the median of the common *ratio*_*thresh*_ range (**Table 2**) or the common *k*_*1,thresh*_ range (**Table 4**). Thus, we were able to validate the threshold values.

### Tumor growth dynamics among stratified populations

We explored the dynamics of the tumor growth for the groups separated by the threshold values to better understand why the anti-VEGF treatment has differential effects in the simulated mouse populations. As researchers have pointed out, log-transformation of tumor growth data provides information on the tumor growth rates (given by the slope of the curve) and is more suitable for detecting a transient biological or therapeutic effect [40, 42, 43]. Therefore, we compared the mean RTV time courses (**Figure S1**) and the log-transformed mean tumor volume data (**Figure S5**) of the groups stratified by the median *ratio*_*thresh*_ (13.869) in each case.

For Roland, Tan, and Volk2008, the mean RTV of the group with larger *k*_*0*_/*k*_*1*_ ratios (**Figure S1A,C,D**) is initially larger, and then becomes smaller relative to the opposite group. This switch occurs because in the group with larger *k*_*0*_/*k*_*1*_ ratios, the difference between the treated and control tumor volumes is smaller at early times, and then becomes larger (**Figure S5**). Meanwhile, the actual tumor volumes for this group are both relatively low. As a result, this group survives longer (Figure 4). For the Mollard case used for validation, the differences between the treated and control tumor volumes in the group with larger *k*_*0*_/*k*_*1*_ ratios are larger (**Figure 6B**, dotted curves), giving rise to the larger mean RTV (**Figure 6A**). However, the group with larger *k*_*0*_/*k*_*1*_ ratios still survives longer because the actual tumor volumes are relatively low (**Figure 5C**).

**Figure 6.**
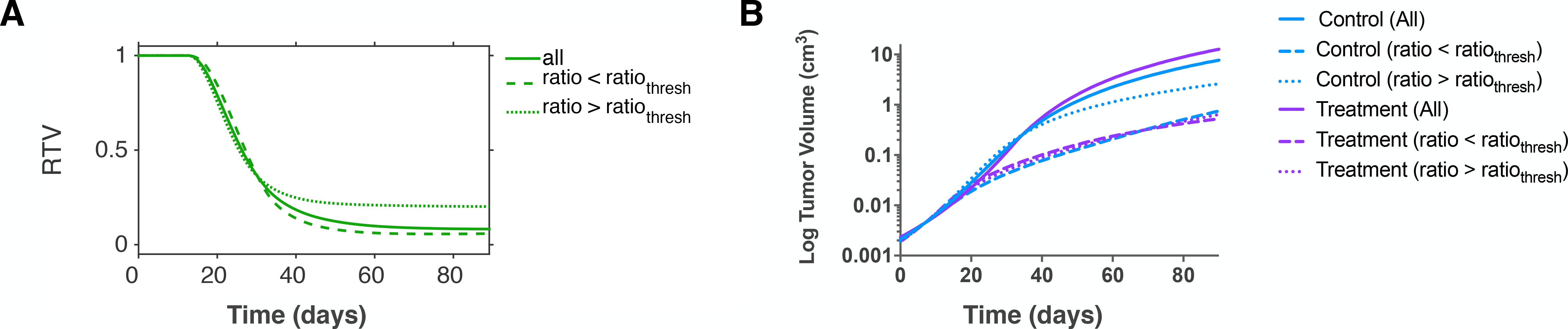
Dynamics of tumor volume. **A**, Time course of relative tumor volume (RTV) for the Mollard case. The mean RTV for all *in silico* mice and mice with tumors whose *k*_*0*_/*k*_*1*_ ratio is smaller and larger than the median *ratio*_*thresh*_ (13.8693) are shown. **B**, Log-transformed tumor volume data for all *in silico* mice and mice separated according to the tumor’s *k*_*0*_/*k*_*1*_ ratio.

The log-transformed tumor volume data also reveal that the tumor growth rates of control and treated groups diverge at different time points. For Roland, the gap between the linear tumor volume data of control and treated groups continually increases (**Figure 7A**). However, the log-transformed tumor volume data show that their growth rates mostly differentiate during days 14-40. The growth rates become similar during the later stage (after 40 days), as evidenced by the parallel curves of the log-transformed data (**Figure 7B**). Therefore, the increasingly large gap between the tumor volumes is a result of early differences in the tumor growth rates. A similar phenomenon is observed for Volk2011b, where the tumor growth rate of the treated group is suppressed transiently at early times but not in the later stage (**Figure S5**). In Zibara, Tan and Volk2008, the growth rates start to differentiate between day 30 and day 45, and only gradually become similar towards the end of the simulated time. Overall, analysis of the log-transformed growth curves reveal that the anti-VEGF treatment has differential effects in limiting tumor growth, and the effects occur at different stages for the simulated cases. The treatment effect appears to be stronger for the group with *k*_*0*_/*k*_*1*_ ratios larger than the median *ratio*_*thresh*_.

**Figure 7.**
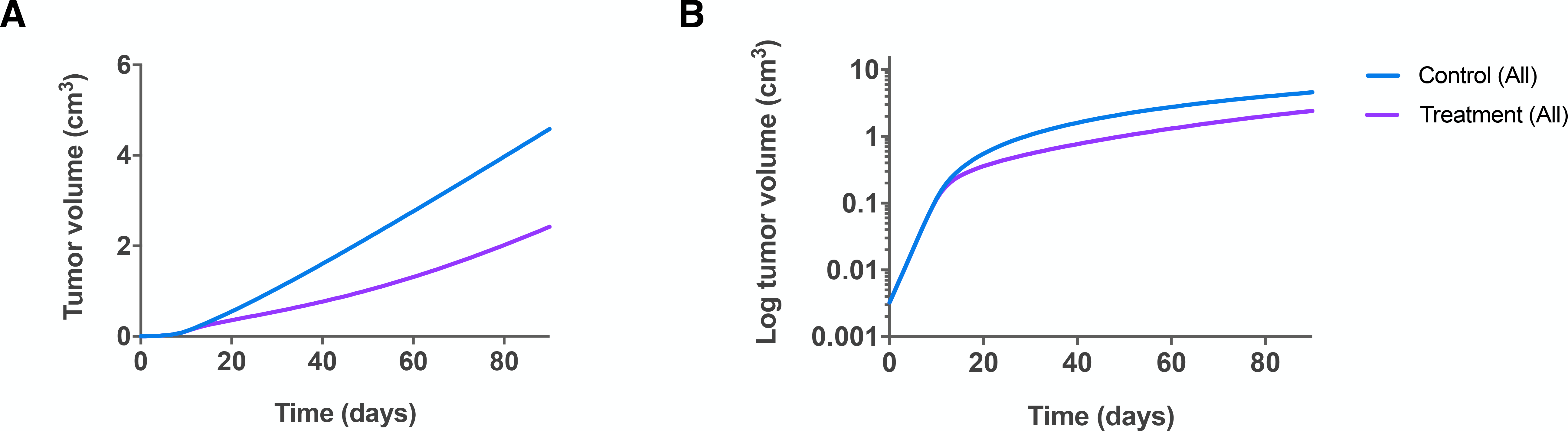
Mean tumor growth. We plot the mean tumor volume for all *in silico* mice in control and treatment groups using the model fitted to Roland data. **A**, Linear scale and **B**, Log scale.

## Discussion

In this study, we focus on identifying potential tumor growth kinetic parameters as biomarkers for the outcome of anti-VEGF treatment. We developed a computational approach to perform biomarker identification that incorporates model training, simulation of tumor growth within a heterogeneous population, and estimation and analysis of population response.

We applied the model to simulate anti-VEGF treatment and compared the effect of treatment across tumor-bearing mice generated from our previous fitting to six independent preclinical studies. For most simulated tumors, the anti-VEGF agent significantly reduces tumor volume compared to control. However, our simulations for Zibara and Volk2011a show that these populations do not respond to the treatment (**Figure 2B,E**), which is different than the effect seen experimentally. This difference occurs for two reasons. First, our simulated treatment protocol A is universal across the six cases, and is different from what was used in each of the original six experimental studies. Second, in our simulations, *k*_*0*_ and *k*_*1*_ are varied simultaneously and independently of each other, possibly resulting in more variability than what occurs in the experimental tumor growth.

Our study demonstrates that the *k*_*0*_/*k*_*1*_ ratio or *k*_*1*_ alone can be utilized to stratify the population response with or without anti-VEGF treatment. This finding agrees with our previous finding through PLSR analysis that the ratio is a key predictor of the tumor response to anti-VEGF treatment [26]. Building on that framework, we found that the survival estimate of mice with larger *k*_*0*_/*k*_*1*_ ratios or smaller *k*_*1*_ is better compared to those with smaller ratios or higher *k_1_.* Interestingly, the result for the ratio is the opposite of the conclusion we drew previously (that a larger ratio correlates with a poorer response to treatment). However, in that work, we focused only on whether the final RTV value was low. This highlights the fact that only evaluating the endpoint RTV of the treated and control group and neglecting the actual tumor volume data over time can lead to misinterpretation of the treatment effect. Indeed, researchers have recognized that while most preclinical studies focus on the end points of tumor growth, monitoring tumor growth kinetically may oftentimes be more insightful [42, 43].

We found that in two cases (Volk2011a simulated with protocol A and protocol V11a), no significant difference is observed in the survival estimates between the pairs of treated and control groups. However, even for these cases, two populations with significantly different survival estimates can be identified based on their *k*_*0*_/*k*_*1*_ ratios (**Figure 4B,E**) or *k*_*1*_ value (**Figure S3B,E**). This indicates that even when the treatment is not effective in reducing tumor volume, there is still a difference in tumor growth dynamics between the two populations stratified based on the tumor’s growth kinetic parameters. Thus, we believe that the *k*_*0*_/*k*_*1*_ ratio or *k*_*1*_ may be prognostic biomarkers to stratify populations for their survival estimate without the anti-angiogenic treatment. Interestingly, the parameters provide mechanistic insight into tumor growth. In particular, they highlight that slower linear growth (larger ratio or smaller *k*_*1*_) results in less aggressive overall tumor growth (**Figure S5**) and therefore, better survival outcome.

Another interesting aspect is the utility of *k*_*1*_ to serve as a prognostic biomarker. Although *k*_*1*_ was not revealed as a strong predictor of the final RTV previously in the PLSR analysis, it is inversely correlated with the *k*_*0*_/*k*_*1*_ ratio, and therefore in our study, it also can be used to stratify the population survival outcome. The use of survival analysis in this study addresses one of the limitations in our previous work. That is, with the PLSR analysis, we were able to identify which parameters were related to treatment efficacy, but could not identify the specific relationship between the kinetic parameter values and effectiveness of the treatment.

Compared to the mean RTV data, the tumor volume data provide more useful insight into the tumor growth characteristics of the stratified population. In particular, the log-transformed tumor volume more clearly illustrates the source of the differences in the population survival estimates. Specifically, we found that larger *k*_*0*_/*k*_*1*_ ratios often yield slower tumor growth in a population, and therefore, lead to a better survival estimate of the population. This conclusion could not be made if we were to only analyze the RTV data. In addition, the log-transformed tumor volume data reveal that the effect of anti-VEGF treatment in tumor growth can be relatively transient (as observed in Roland and Volk2011b) or gradual (as seen in Zibara, Tan, and Volk2008).

Our study makes use of a predictive and useful computational model of tumor growth with and without anti-VEGF treatment. This is a pharmacokinetics-pharmacodynamics model with mechanistic detail that goes beyond what is found in other models. However, in the future, this model can be expanded to address limitations that are not currently accounted for. For example, we do not account for changes in tumor vascularity relative to tumor volume. In addition, vascular normalization is an important process that has been shown to affect tumor growth and can be regulated by anti-VEGF agents [32]; however, this process is not included in our model. These aspects can be implemented into the model as more quantitative data become available and enable us to characterize the dynamics of vessel normalization. The model can then be further extended to account for other characteristics of tumor progression, including tumor perfusion and metastatic potential. Furthermore, the range of threshold values for tumor stratification is constrained to be within the overlap of estimated parameter values from model training to each experimental dataset. It is possible that artifacts coming from experimental data quantification led to bias in the range of the fitted parameter values. This can also be improved when more quantitative data from experimental studies become available to allow for additional model training, which enables improved model predictions. We note that the biomarker candidates identified in this study are best used to stratify populations for their survival outcome, whether the mice receive treatment or not, rather than to predict treatment efficacy. This utility of the growth parameters for stratification is primarily because the datasets used to train the model were the control time courses of tumor growth over several weeks. Our results would be of broader applicability if only pre-treatment data were adequate to train the model. We have attempted such approach in our previous study [26]; however, the fitting results were not satisfactory (predicted volumes varied widely, preventing us from making conclusive predictions). Despite this perceived limitation, our modeling approach generates hypotheses about potential biomarkers, and further experimental validation is needed to ensure the utility of the biomarkers identified here.

Our study demonstrates a time- and cost-effective way to generate large *in silico* mouse populations, predict anti-VEGF treatment outcome, and stratify the populations. This approach provides useful information that could facilitate efficient experimental design, such as predicting the effect of different treatment protocols (varying the dosage and the timing of the injections). Additionally, our modeling approach can be adapted for analysis of patient treatment outcome in clinical studies. With data from a small patient population, we can develop a model that is trained to patient-specific data and generate a larger *in silico* patient population. Analysis of the simulated tumor growth and survival data can be used to identify biomarkers that predict responders versus non-responders to anti-VEGF treatment, stratify the predicted population survival, and test the response to various treatment schedules.

## Conclusion

We examined tumor growth kinetic parameters as potential biomarkers of anti-angiogenic treatment outcome. Using a computational model that simulates VEGF-dependent tumor growth in tumor-bearing mice, we generated an *in silico* mouse population and related the kinetic parameters that characterize tumor growth to the response to anti-VEGF treatment. We found that the ratio between two tumor growth kinetic parameters, *k*_*0*_ and *k*_*1*_, as well as *k*_*1*_ alone, can be prognostic biomarkers and that the simulated treatment protocol may have a better outcome for mice whose tumors have smaller linear growth rates. In fact, we found ranges of threshold values for the *k*_*0*_/*k*_*1*_ ratio and *k*_*1*_ that distinguish tumors’ response to the anti-VEGF treatment. This study demonstrates an approach for identifying tumor growth kinetic parameters as potential biomarkers, and this model framework can be adapted to predict the efficacy of other anti-angiogenic strategies.

## Methods

### Computational model

This work directly uses our previously calibrated and validated three-compartment model of a tumor-bearing mouse [26]. In this previous work, we fit the model to six independent control datasets, and the model-predicted tumor growth curves match closely to the experimental data (fitting error range: 0.0405 − 0.1833). We provide a full description of this model in **Supplemental File S1**.

### Numerical implementation

All model equations were implemented in MATLAB using the SimBiology toolbox. The model is provided as the SimBiology project file, as SBML, and as a MATLAB m-file (**Supplemental File S2**). Parameter fitting was performed using the *Isqnonlin* function in MATLAB. Kaplan-Meier survival estimation was performed using the *kmplot* function in MATLAB, and GraphPad Prism was used for statistical survival analyses.

### Simulation of *in silico* mouse population

In our previous work, we fit the three-compartment model of a tumor-bearing mouse to six datasets. Here, we generated 400 sets of values for growth parameters *k*_*0*_ and *k*_*1*_, randomly selected from a uniform distribution within the range of the best-fit parameter sets from our previous study (**Table S1**). The *Ang*_*0*_ value is set to be the median of the best fits in each case (**Table S1**). These sets were used to calculate tumor growth with or without anti-VEGF treatment, simulating a population of mice for each of the six datasets. In order to keep tumor growth profiles realistic, tumors that do not reach 0.1 cm^3^ within 10 days upon tumor engraftment (assuming tumor volume to be 0.004 cm^3^) were excluded from the analyses.

For each dataset, we simulated anti-VEGF treatment as intravenous injections lasting for one minute. We implement this injection by adding a net rate of secretion of the drug (bevacizumab, which binds to human VEGF isoforms) directly into the blood compartment (**Figure 1**). We simulated different anti-VEGF treatment protocols. Treatment protocol A is simulated universally across the six cases. In this protocol, weekly treatment starts when the tumor volume reached 0. 1 cm^3^, as the switch where angiogenesis is more strongly promoted occurs when the tumor reaches 1-2 mm in diameter. The treatment dosage is 10 mg/kg. The model was simulated for 12 weeks after treatment started. We also simulated alternate treatment protocols: Z, denotes biweekly treatment at dosage of 10 mg/kg starting when tumor volume is 0.004 cm^3^; V11a, denotes biweekly treatment (twice a week) at a dosage of 10 mg/kg, starting when the tumor volume is 0.5 cm^3^; and V11a-D, denotes biweekly treatment at a dosage of 20 mg/kg, starting when tumor volume is 0.5 cm^3^. Information for all treatment protocols is shown in **Table S3**.

### Relative tumor volume (RTV)

Based on the model-generated tumor growth data, the relative tumor volume (RTV), the ratio between the treated and control tumor volumes, is calculated at any simulated time point:

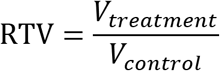

An RTV value less than one indicates that the treated tumor volume is smaller than the control.

### Kaplan-Meier survival estimation

We applied time-to-event analysis to determine the survival of each mouse population [34]. An *in silico* mouse is recorded as “sacrificed” when its tumor reaches 2 cm^3^ within the simulated time. Alternatively, a mouse is recorded as “censored” at a particular time point, *t*, if its tumor volume simulation remains below 2 cm^3^ but ended before that time *t*. All other mice are retained in the study and recorded as “alive”. Survival curves were estimated by the Kaplan-Meier method using the *kmplot* function in MATLAB [44], and compared using the Mantel-Cox log rank test and Mantel-Haenszel hazard ratio in GraphPad Prism.

The hazard ratio (HR) compares the rate of death in two groups, with the assumption that the population hazard ratio is consistent over time. It is calculated using the Mantel-Haneszel approach, which is more accurate than the log rank approach [45]. As an example, an HR of 0.5 between two groups means that the death rate of the first group is half of that of the second group.

### Determination of threshold values

In order to determine threshold values for the *k*_*0*_/*k*_1_ ratio, we ordered the simulated mouse tumor volume data for each of the six populations according to the *k*_*0*_/*k*_1_ ratio. Then, we systematically tested each *k*_*0*_/*k*_*1*_ ratio (called “*ratio*_*thresh*_”) value to see if there is a significant difference between the survival estimates for the mice with *k*_*0*_/*k*_*1*_ ratio above and below “*ratio*_*thresh*_” in the log rank test (*p*<0.05). We performed a similar analysis for *k*_*0*_ and *k*_*1*_ individually to determine any *k*_*0,thresh*_ and *k*_*1,thresh*_ values.

### Validation of the predicted biomarker

Upon identifying a potential predictive biomarker for the efficacy of anti-VEGF treatment, we validated our findings using an independent set of data that was not used to determine the range of the threshold value. To do so, we fit the control tumor growth for the independent data set and generated an *in silico* mouse population based on the fitted parameters.

#### Data extraction

For threshold validation, data from the published *in vivo* experimental study of MDA-MB-231 xenograft tumor growth in mice by Mollard *et al.* were used for parameter estimation [41]. The dataset for tumor growth in untreated tumors (control) was used for fitting the model, and the dataset for tumors treated with bevacizumab twice weekly was used for validation. Experimental data was extracted using the WebPlotDigitizer program [46]. The numerical values are provided in **Table S1**.

#### Parameter estimation

We trained the model to fit the control tumor growth dataset from [41] using the same approach as described in our previous work [26]. The values of tumor growth parameters *k*_*0*_, *k*_*1*_, and *Ang*_*0*_ were estimated. In their study, Mollard and coworkers only reported the tumor volumes relative to day eight. However, the absolute tumor volumes are needed to determine how the tumor interstitial volume varies as a function of the total tumor volume. Therefore, we compared the relative tumor volume at each time point in the work by Mollard and coworkers to that of all the available control datasets (**Figure S6**). We then chose to use the interstitial volume equation from the Zibara data, given that the relative tumor volume closely matches that of the data in Mollard. Finally, we fit our tumor growth model to the Mollard control dataset using the *lsqnonlin* function in MATLAB to minimize the sum of squared residuals:

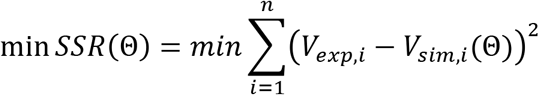

where *V*_*exp,I*_ is the *i*th experimental data point of tumor volume, *V*_*sim,I*_ is the *i*th simulated volume at the corresponding time point, and *n* is the total number of experimental data points. The minimization is subject to Θ, the set of upper and lower bounds on each of the free parameters.

The bounds for each parameter spanned at least one order of magnitude: 10^−8^ to 10^−2^ for *k*_*0*_ and *k*_*1*_ and 10^−16^ to 10^−14^ for *Ang_0_*. After fitting to the control data, we validated the estimated parameters with measured tumor volumes not used in the fitting for model validation: the experimentally measured volumes of tumors treated with the anti-angiogenic drug bevacizumab. We simulated the dosing regimen used in the experiment performed by Mollard *et al.* using the estimated parameters obtained from fitting to the control data. This protocol involved three cycles of weekly intravenous injections lasting for one minute starting from day five. We used the combined SSR for the relative tumor volume between model prediction and the experimental data (both control and treatment) to identify the optimal parameters. Twelve parameter sets with the smallest errors were taken to be the “best” sets (**Table S1**) and the ranges of the estimated parameter values were used for subsequent model simulations (**Table S2**).

We extracted the absolute tumor volume at day 8 from previously reported data from Mollard and coworkers [47] to determine the survival estimates for a mouse population simulated based on the fitted growth kinetics parameter values.

## Declarations

### Ethics approval and consent to participate

Not applicable

### Consent for publication

Not applicable

### Data accessibility

All data analyzed during this study, including the model used for data generation, are included in this published article and its supplementary information files.

### Author’s contributions

SDF designed the research. ADA wrote MATLAB scripts and performed initial model simulations. QW performed model simulations and data analysis. QW and SDF wrote the manuscript. All authors have read and approved the final manuscript, and agree to be accountable for all aspects of the work.

## Acknowledgements

The authors thank members of the Finley research group for critical comments and suggestions.

## Supplemental Figures

**Figure S1.** Time course of relative tumor volume (RTV). The mean RTV levels of all *in silico* mice and the groups with *k*_*0*_/*k*_*1*_ smaller or larger than the median *ratio*_*thresh*_ (13.8693) are shown. **A**, Roland. **B**, Zibara. **C**, Tan. **D**, Volk2008. **E**, Volk2011a. **F**, Volk2011b.

**Figure S2.** Scatter plot of RTV at the end of simulations versus tumor growth kinetic parameters. Left to right columns: *k*_*0*_, *k*_*1*_, and *k*_*0*_/*k*_*1*_ ratio. **A**, Roland. **B**, Zibara. **C**, Tan. **D**, Volk2008. **E**, Volk2011a. **F**, Volk2011b. Color gradient represents the range of RTV values (from 0 to 1).

**Figure S3.** Kaplan-Meier curves for the six simulated groups of tumor-bearing mice. Here, the *k*_*1,thresh*_ value is taken as the median from the common range found among the six cases (1.661 =10^−6^). **A**, Roland. **B**, Zibara. **C**, Tan. **D**, Volk2008. **E**, Volk2011a. **F**, Volk2011b. The estimated survival curves of *in silico* mice subgroups within each group are shown in each plot: all mice, mice with *k*_*1*_ smaller or larger than the median *k*_*1,thresh*_, in the control setting or with treatment.

**Figure S4.** Model-simulated tumor growth data with alternative treatment protocols. The mean and 95% confidence interval at each time point are shown. **A**, Zibara case with treatment protocol Z. **B**, Volk2011a case with treatment protocol V11a. **C**, Volk2011a case with treatment protocol V11a-D (see Methods). Asterisks indicate that the difference between the control group and the treatment group tumor volumes is statistically significant (*p*<0.05).

**Figure S5.** Log-transformed tumor volume data of all *in silico* mice and populations separated by *ratio*_*thresh*_ value of 13.8693. **A**, Roland. **B**, Zibara. **C**, Tan. **D**, Volk2008. **E**, Volk2011a. **F**, Volk2011b.

**Figure S6.** Comparison of normalized experimental data. Control tumor volume on day eight is extrapolated from an exponential fit to the experimental data [35–39], and used to calculate the relative tumor volumes for the models fit to the control tumor volume from each of the six datasets. The resulting normalized control tumor volume datasets are compared to that from the Mollard study.

## Supplementary Material

### Supplemental Tables

Table S1: Experimental measurements and estimated parameter values from fitting to Mollard dataset

Table S2: Parameter bounds and values used in simulations

Table S3: Description of treatment protocols

### Supplemental Figures

Figure S1. Time course for the relative tumor volume (RTV) vs. time

Figure S2. Relationship between RTV and tumor growth kinetic parameters

Figure S3. Kaplan-Meier curves for tumor-bearing mouse population separated using *k_1, thresh_*

Figure S4. Tumor volume time course for Z and V11a, V11a-D protocols

Figure S5. Time course for log-transformed tumor volume

Figure S6. Experimental data for normalized control tumor volume extracted from published datasets

### Supplemental Files

File S1. Description of three-compartment model

File S2. Zipped file containing the computational model, as a MATLAB SimBiology file, SBML, and a MATLAB .m file

File S3. Supplemental Tables

File S4. Supplemental Figures

## References

1. Carmeliet P, Jain RK. 2011 Molecular mechanisms and clinical applications of angiogenesis. Nature 473:298–307 (doi:10.1038/nature10144).

2. Belal A, Maha A, Morgan T, L. dd, R. sp. 2012 Antiangiogenic therapy for cancer: an update. Pharmacotherapy 32:1095–111 (doi:10.1002/phar.1147).

3. FDA Approval for Bevacizumab. National Cancer Institute https://www.cancer.gov/about-cancer/treatment/drugs/fda-bevacizumab. Accessed 2 Apr 2018.

4. Montero AJ, Escobar M, Lopes G, Glück S, Vogel C. 2012 Bevacizumab in the treatment of metastatic breast cancer: friend or foe? Curr Oncol Rep 14:1–11 (doi:10.1007/s11912-011-0202-z).

5. Lambrechts D, Lenz H-J, de Haas S, Carmeliet P, Scherer SJ. 2013 Markers of response for the antiangiogenic agent bevacizumab. J Clin Oncol 31:1219–30 (doi:10.1200/JCO.2012.46.2762).

6. Rubovszky G, Horváth Z. 2017 Recent advances in the neoadjuvant treatment of breast cancer. J Breast Cancer 20:119–31 (doi:10.4048/jbc.2017.20.2.119).

7. Carey LA, Berry DA, Cirrincione CT, Barry WT, Pitcher BN, Harris LN, et al. 2016 Molecular heterogeneity and response to neoadjuvant human epidermal growth factor receptor 2 targeting in calgb 40601, a randomized phase iii trial of paclitaxel plus trastuzumab with or without lapatinib. J Clin Oncol 34:542–9 (doi:10.1200/JCO.2015.62.1268).

8. Untch M, Loibl S, Bischoff J, Eidtmann H, Kaufmann M, Blohmer J-U, et al. 2012 Lapatinib versus trastuzumab in combination with neoadjuvant anthracycline-taxane-based chemotherapy (GeparQuinto, GBG 44): a randomised phase 3 trial. Lancet Oncol 13:135–44 (doi:10.1016/S1470-2045(11)70397-7).

9. Bear HD, Tang G, Rastogi P, Geyer CE, Liu Q, Robidoux A, et al. 2015 Neoadjuvant plus adjuvant bevacizumab in early breast cancer (NSABP B-40 [NRG Oncology]): secondary outcomes of a phase 3, randomised controlled trial. Lancet Oncol 16:1037–48 (doi:10.1016/S1470-2045(15)00041-8).

10. Earl HM, Hiller L, Dunn JA, Blenkinsop C, Grybowicz L, Vallier A-L, et al. 2015 Efficacy of neoadjuvant bevacizumab added to docetaxel followed by fluorouracil, epirubicin, and cyclophosphamide, for women with HER2-negative early breast cancer (ARTemis): an open-label, randomised, phase 3 trial. Lancet Oncol 16:656–66 (doi:10.1016/S1470-2045(15)70137-3).

11. Nahleh ZA, Barlow WE, Hayes DF, Schott AF, Gralow JR, Sikov WM, et al. 2016 SWOG S0800 (NCI CDR0000636131): addition of bevacizumab to neoadjuvant nab-paclitaxel with dose-dense doxorubicin and cyclophosphamide improves pathologic complete response (pCR) rates in inflammatory or locally advanced breast cancer. Breast Cancer Res Treat 158:485–95 (doi:10.1007/s10549-016-3889-6).

12. Rugo HS, Olopade OI, DeMichele A, Yau C, van’t Veer LJ, Buxton MB, et al. 2016 Adaptive randomization of veliparib-carboplatin treatment in breast cancer. N Engl J Med 375:23–34 (doi:10.1056/NEJMoa1513749).

13. Wehland M, Bauer J, Magnusson NE, Infanger M, Grimm D. 2013 Biomarkers for anti-angiogenic therapy in cancer. Int J Mol Sci 14:9338–64 (doi:10.3390/ijms14059338).

14. Burstein HJ, Chen Y-H, Parker LM, Savoie J, Younger J, Kuter I, et al. 2008 Vegf as a marker for outcome among advanced breast cancer patients receiving anti-vegf therapy with bevacizumab and vinorelbine chemotherapy. Clin Cancer Res 14:7871–7 (doi:10.1158/1078-0432.CCR-08-0593).

15. Jain RK, Duda DG, Willett CG, Sahani DV, Zhu AX, Loeffler JS, et al. 2009 Biomarkers of response and resistance to antiangiogenic therapy. Nat Rev Clin Oncol 6:327–38 (doi:10.1038/nrclinonc.2009.63).

16. Willett CG, Boucher Y, di Tomaso E, Duda DG, Munn LL, Tong RT, et al. 2004 Direct evidence that the VEGF-specific antibody bevacizumab has antivascular effects in human rectal cancer. Nat Med 10:145–7 (doi:10.1038/nm988).

17. Willett CG, Boucher Y, Duda DG, di Tomaso E, Munn LL, Tong RT, et al. 2005 Surrogate markers for antiangiogenic therapy and dose-limiting toxicities for bevacizumab with radiation and chemotherapy: continued experience of a phase I trial in rectal cancer patients. J Clin Oncol 23:8136–9 (doi:10.1200/JCO.2005.02.5635).

18. Willett CG, Duda DG, di Tomaso E, Boucher Y, Ancukiewicz M, Sahani DV, et al. 2009 Efficacy, safety, and biomarkers of neoadjuvant bevacizumab, radiation therapy, and fluorouracil in rectal cancer: a multidisciplinary phase II study. J Clin Oncol 27:3020–6 (doi:10.1200/JCO.2008.21.1771).

19. Zhu AX, Sahani DV, Duda DG, di Tomaso E, Ancukiewicz M, Catalano OA, et al. 2009 Efficacy, safety, and potential biomarkers of sunitinib monotherapy in advanced hepatocellular carcinoma: a phase ii study. J Clin Oncol 27:3027–35 (doi:10.1200/JCO.2008.20.9908).

20. Batchelor TT, Duda DG, di Tomaso E, Ancukiewicz M, Plotkin SR, Gerstner E, et al. 2010 Phase II study of cediranib, an oral pan-vascular endothelial growth factor receptor tyrosine kinase inhibitor, in patients with recurrent glioblastoma. J Clin Oncol 28:2817–23 (doi:10.1200/JCO.2009.26.3988).

21. Lee JH, Lee HY, Ahn M-J, Park K, Ahn JS, Sun J-M, et al. 2016 Volume-based growth tumor kinetics as a prognostic biomarker for patients with EGFR mutant lung adenocarcinoma undergoing EGFR tyrosine kinase inhibitor therapy: a case control study. Cancer imaging 16 (doi:10.1186/s40644-016-0063-7).

22. Seyal AR, Parekh K, Arslanoglu A, Gonzalez-Guindalini FD, Tochetto SM, Velichko YS, et al. 2015 Performance of tumor growth kinetics as an imaging biomarker for response assessment in colorectal liver metastases: correlation with FDG PET. Abdom imaging 40:3043–51.

23. El Sharouni S, Kal H, Battermann J. 2003 Accelerated regrowth of non-small-cell lung tumours after induction chemotherapy. Br J Cancer (doi:10.1038/sj.bjc.6601418).

24. Stein WD, Yang J, Bates SE, Fojo T. 2008 Bevacizumab reduces the growth rate constants of renal carcinomas: a novel algorithm suggests early discontinuation of bevacizumab resulted in a lack of survival advantage. Oncologist 13:1055–62 (doi:10.1634/theoncologist.2008-0016).

25. Rezai P, Yaghmai V, Tochetto S, Galizia M, Miller F, Mulcahy MF, et al. 2011 Change in the growth rate of localized pancreatic adenocarcinoma in response to gemcitabine, bevacizumab, and radiation therapy on MDCT. Int J Radiat Oncol Biol Phys 81:452–9 (doi:10.1016/j.ijrobp.2010.05.060).

26. Gaddy TD, Wu Q, Arnheim AD, Finley SD. 2017 Mechanistic modeling quantifies the influence of tumor growth kinetics on the response to anti-angiogenic treatment. PLoS Comput Biol 13:e1005874 (doi:10.1371/journal.pcbi.1005874).

27. Martin EC, Aarons L, Yates JWT. 2016 Accounting for dropout in xenografted tumour efficacy studies: integrated endpoint analysis, reduced bias and better use of animals. Cancer Chemother Pharmacol 78:131–41 (doi:10.1007/s00280-016-3059-x).

28. Claret L, Bruno R. 2014 Assessment of tumor growth inhibition metrics to predict overall survival. Clin Pharmacol Ther 96:135–7 (doi:10.1038/clpt.2014.112).

29. Altrock PM, Liu LL, Michor F. 2015 The mathematics of cancer: integrating quantitative models. Nat Rev Cancer 15:730–45 (doi:10.1038/nrc4029).

30. Masoudi-Nejad A, Wang E. 2015 Cancer modeling and network biology: Accelerating toward personalized medicine. Seminars in Cancer Biology 30:1–3 (doi:10.1016/j.semcancer.2014.06.005).

31. Yankeelov TE, An G, Saut O, Luebeck EG, Popel AS, Ribba B, et al. 2016 Multi-scale modeling in clinical oncology: opportunities and barriers to success. Ann Biomed Eng 44:2626–41 (doi:10.1007/s10439-016-1691-6).

32. Jain RK. 2013 Normalizing tumor microenvironment to treat cancer: bench to bedside to biomarkers. J Clin Oncol 31:2205–18 (doi:10.1200/JCO.2012.46.3653).

33. Jubb AM, Miller KD, Rugo HS, Harris AL, Chen D, Reimann JD, et al. 2011 Impact of exploratory biomarkers on the treatment effect of bevacizumab in metastatic breast cancer. Clin Cancer Res 17:372–81 (doi:10.1158/1078-0432.CCR-10-1791).

34. Bender BC, Schindler E, Friberg LE. 2015 Population pharmacokinetic-pharmacodynamic modelling in oncology: a tool for predicting clinical response. Br J Clin Pharmacol 79:56–71 (doi:10.1111/bcp.12258).

35. Roland CL, Dineen SP, Lynn KD, Sullivan LA, Dellinger MT, Sadegh L, et al. 2009 Inhibition of vascular endothelial growth factor reduces angiogenesis and modulates immune cell infiltration of orthotopic breast cancer xenografts. Mol Cancer Ther 8:1761–71 (doi:10.1158/1535-7163.MCT-09-0280).

36. Zibara K, Awada Z, Dib L, El-Saghir J, Al-Ghadban S, Ibrik A, et al. 2015 Anti-angiogenesis therapy and gap junction inhibition reduce MDA-MB-231 breast cancer cell invasion and metastasis in vitro and in vivo., Anti-angiogenesis therapy and gap junction inhibition reduce MDA-MB-231 breast cancer cell invasion and metastasis in vitro and in vivo. Sci Rep 5, 5:12598–12598 (doi:10.1038/srep12598, 10.1038/srep12598).

37. Tan G, Kasuya H, Tevfik Tolga Sahin, Kazuo Yamamura, Zhiwen Wu, Yusuke Koide, et al. 2014 Combination therapy of oncolytic herpes simplex virus HF10 and bevacizumab against experimental model of human breast carcinoma xenograft. Int J Cancer 136:1718–30 (doi:10.1002/ijc.29163).

38. Volk LD, Flister MJ, Bivens CM, Stutzman A, Desai N, Trieu V, et al. 2008 Nab-paclitaxel efficacy in the orthotopic model of human breast cancer Is significantly enhanced by concurrent anti-vascular endothelial growth factor A therapy. Neoplasia 10:613–23 (doi:10.1593/neo.08302).

39. Volk LD, Flister MJ, Chihade D, Desai N, Trieu V, Ran S. 2011 Synergy of nab-paclitaxel and bevacizumab in eradicating large orthotopic breast tumors and preexisting metastases. Neoplasia 13:327–38 (doi:10.1593/neo.101490).

40. Hather G, Liu R, Bandi S, Mettetal J, Manfredi M, Shyu W-C, et al. 2014 Growth rate analysis and efficient experimental design for tumor xenograft studies. Cancer Inform 13 Suppl 4:65–72 (doi:10.4137/CIN.S13974).

41. Mollard S, Ciccolini J, Imbs D-C, Cheikh RE, Barbolosi D, Benzekry S. 2017 Model driven optimization of antiangiogenics + cytotoxics combination: application to breast cancer mice treated with bevacizumab + paclitaxel doublet leads to reduced tumor growth and fewer metastasis. Oncotarget 8:23087–98 (doi:10.18632/oncotarget.15484).

42. Gengenbacher N, Singhal M, Augustin HG. 2017 Preclinical mouse solid tumour models: status quo, challenges and perspectives. Nat Rev Cancer 17:751–65 (doi:10.1038/nrc.2017.92).

43. Nasarre P, Thomas M, Kruse K, Helfrich I, Wolter V, Deppermann C, et al. 2009 Host-derived angiopoietin-2 affects early stages of tumor development and vessel maturation but is dispensable for later stages of tumor growth. Cancer Res 69:1324–33 (doi:10.1158/0008-5472.CAN-08-3030).

44. KMPlot - File Exchange - MATLAB Central. http://www.mathworks.com/matlabcentral/fileexchange/22293-kmplot. Accessed 2 Apr 2018.

45. GraphPad Statistics Guide. https://www.graphpad.com/guides/prism/7/statistics/index.htm?stat_howto_survival.htm. Accessed 2 Apr 2018.

46. WebPlotDigitizer - Extract data from plots, images, and maps. https://automeris.io/WebPlotDigitizer/. Accessed 2 Apr 2018.

47. Mollard S, Fanciullino R, Giacometti S, Serdjebi C, Benzekry S, Ciccolini J. 2016 In vivo bioiluminescence tomography for monitoring breast tumor growth and metastatic spreading: comparative study and mathematical modeling. Sci Rep 6:36173 (doi:10.1038/srep36173).

